# Genomic evaluation of *Bordetella spp*. originating from Australia

**DOI:** 10.1101/2021.03.02.433639

**Authors:** Winkie Fong, Verlaine Timms, Eby Sim, Vitali Sintchenko

## Abstract

*Bordetella pertussis* is the primary causative agent of pertussis, a highly infectious respiratory disease associated with prolonged coughing episodes. Pertussis infections are typically mild in adults, however in neonates, infections can be fatal. Despite successful vaccine uptake, the disease is re-emerging across the globe, therefore it is critical to determine the mechanism by which *B. pertussis* is escaping vaccination control. Studies have suggested that significant changes have occurred in *B. pertussis* genomes in response to whole cell and acellular vaccines. Continued molecular monitoring is therefore crucial for public health surveillance.

High-resolution molecular surveillance of *B. pertussis* can be achieved through the sequencing of the whole genome. In public health laboratories, whole genome sequencing is primarily performed by short-read sequencing technologies as they are most cost-effective. However short read sequencing does not resolve the extensive genomic rearrangement evident in *Bordetella* genomes. This is because repeat regions present in *Bordetella* genomes are collapsed by downstream analysis. For example, the *B. pertussis* genome contains more than 200 copies of the IS*481* insertion element, hence assemblies generally consist of >200 contigs. Advancements in long-read technologies however increase the potential to circularise and close genomes by bridging the locations of the IS*481* insertion element.

In this study, we aimed to contextualise the *Bordetella* spp. circulating in NSW, Australia and assess their relationship with global isolates utilising core genome, SNP and structural clustering analysis using long read technology. We report five closed genomes of *Bordetella* spp. isolated from Australian patients. Two of the three *B. pertussis* closed isolates, were unique with their own genomic structure, while the other structurally clustered with global isolates. We found that Australian *B. holmesii* and *B. parapertussis* strains cluster with global isolates and do not appear to be unique to Australia. Australian draft *B. holmesii* SNP analysis showed that between 1999 and 2007, isolates were relatively similar, however post-2012, isolates were distinct from each other. The closed isolates can also be used as high-quality reference sequences for both surveillance and other investigations into pertussis spread.

## Background

Pertussis, or whooping cough, caused by *Bordetella pertussis* (and occasionally *Bordetella parapertussis* and *Bordetella holmesii*) is a severe respiratory infection characterized by prolonged coughing episodes^1^. In the pre-vaccination era, pertussis was the foremost cause of infantile death due to an infectious disease in the first year of life^1^. Recently however, pertussis has been re-emerging over the past few decades and this is despite high vaccination coverage rates. The causes of this resurgence are possibly related to vaccine escape, waning immunity or poor vaccine compliance. Further, with the rise of molecular based diagnostic tests it is now clear that other *Bordetella* species such as *Bordetella parapertussis* and *Bordetella holmesii* can also cause a mild form of pertussis-like illness. In Australia, *B. holmesii* can circulate at a prevalence of between 6.5 - 16.8%^2^ and is indistinguishable from *B. pertussis* when using the common PCR diagnostic target of *IS481*^3^.

Another possible factor contributing to the re-emergence of *B. pertussis* is a change in the antigenic structure of circulating strains. Current acellular vaccines vary between countries and contain 2-5 immunogenic antigens produced by *B. pertussis* – Pertussis toxin (PT), Filamentous haemagglutinin (FHA), Pertactin (PRN), Fimbrae (FIM) and the adenylate cyclase toxin (ACT). This has potentially led to the natural selection of circulating strains with antigenic variants not present in the vaccine. Changes in antigenic variation have been documented independently in several countries, including Australia, such as the emergence of PRN-deficient^4, 5^, FHA-deficient^6^ or PT-deficient^7^ strains of *B. pertussis*. It also appears that over recent years, *B. pertussis* strains have diverged from historical clones to produce a novel allele for the PT promoter *ptxP* ^*8*^. Prior to the introduction of the vaccine, circulating strains encoded *ptxP1*, however, current predominating strains contain *ptxP3*. Furthermore, these recent *ptxP3* strains also produce 1.5 times more PT than their *ptxP1* counterparts^8^.

Molecular surveillance of respiratory pathogens such as *B. pertussis* is paramount for effective disease control^9^. Molecular typing of *B. pertussis* has been transformed with the advent of higher-resolution genome sequencing methods such as whole genome sequencing (WGS). Available isolates have allowed the development of single nucleotide polymorphism (SNP) typing that is WGS-based. *B. pertussis* strains can be classified into SNP clusters (I-VI)^10^. SNP clusters can be defined by specific SNP changes in genes, the primary SNP change being in *ptxP* (G > A mutation in the intergenic region of *B. pertussis* Tohama I BP3782 - designated *ptxP3*) and thus the sole definer of SNP Cluster I^10^. In Australia, SNP Cluster I can be further divided into 5 epidemic lineages (EL1-EL5)^11^.

WGS can provide a wealth of information that can be utilised for SNP-based typing or observations of the vaccine antigen sequences using *de novo* assembly methods. However, short-read based *de novo* assembly algorithms conglomerate repeat regions of the genome resulting in basic and limited understandings of genomic structure and complexity^12^. This is problematic in the assembly of the *B. pertussis* genome since it contains more than 200 copies of the IS*481* insertion element^13^. As a result, short-read assemblies are highly fragmented, (often >200 contigs) which can lead to information being missed. Previous research has also determined that the *B. pertussis* genome is subject to large genomic inversions and rearrangements, with most mediated by *IS481*^*14*^. The type and positions of the rearrangements are grouped by *B. pertussis* phylogenetic clusters^14^. However, these structural rearrangements are not unique to *B. pertussis*, in fact, *B. parapertussis* and *B. holmesii* also carry these characteristics^15^.

Generating high-quality reference sequences that represent locally circulating strains can more accurately identify variable regions of the genome, virulence and antibiotic resistance markers and greatly support the accuracy of phylogenetic analysis^16, 17^. Advancements in long-read technologies like Nanopore and PacBio sequencing have increased the potential to circularise and close genomes by bridging the locations of the *IS481* insertion element^18-20^. Further, as these genomic rearrangements have only been investigated in strains from the northern hemisphere, primarily the US, the genomic rearrangements prevalent in strains circulating in Australia and their significance for molecular surveillance is currently unknown.

Therefore, this study sought to investigate the mechanisms of the potential evolution of *Bordetella* spp. in Australia using a combination of long and short read technologies to contextualise genomic rearrangements within previously defined SNP lineages. The study present five closed *Bordetella* spp. genomes from the southern hemisphere that can be used as references in future studies investigating *Bordetella* spp. strains circulating in Australia.

## Results

### Sequencing Statistics

Five isolates were sequenced using both Nanopore long read sequencing and Illumina short reads to close and polish the genome. Statistics of the sequencing results are presented in Table 1.

**Table 1:** Summary statistics of closed *Bordetella* spp. genomes sequenced

| Sample | Total Reads | Mean Read Length | Read Length N50 | Genome Length | GC content (%) | Count of CDS | Count of tRNAs | Count of rRNA |
| --- | --- | --- | --- | --- | --- | --- | --- | --- |
| CIDM-BH3 | 467,478 | 4,731.3 | 9,416 | 3,696,928 | 62.69 | 3,683 | 56 | 9 |
| CIDM-BP2 | 501,426 | 6,830.1 | 14,169 | 4,109,518 | 67.70 | 3,971 | 64 | 9 |
| CIDM-BP3 | 622,884 | 8,636.8 | 17,331 | 4,093,865 | 67.71 | 3,954 | 64 | 9 |
| CIDM-BP5† | 41,276 | 8,813 | 16,164 | 4,104,145 | 67.71 | 3,943 | 64 | 9 |
| CIDM-BPP2 | 138,446 | 6,830.1 | 20,730 | 4,775,256 | 68.10 | 4,573 | 65 | 9 |

†Sequencing was only ran for 24 hours

In addition, we further sequenced seven *B. holmesii* isolates using short read technology and sequencing and assembly statistics are listed in Table 2.

**Table 2:** Summary of the Illumina sequencing and assembly statistics of all *B. holmesii* isolates.

| Sample | Total Reads | N50 | Contigs (>1000 bp) | Genome Coverage (%) | GC content (%) | Count of CDS |
| --- | --- | --- | --- | --- | --- | --- |
| CIDM-BH1 | 3,614,054 | 29,344 | 189 | 94.55 | 62.76 | 3,396 |
| CIDM-BH2 | 3,432,966 | 29,426 | 188 | 93.87 | 62.79 | 3,557 |
| CIDM-BH4 | 2,588,521 | 26,580 | 197 | 93.87 | 62.78 | 3,383 |
| CIDM-BH6 | 2,542,224 | 27,377 | 197 | 93.91 | 62.78 | 3,386 |
| CIDM-BH7 | 2,564,461 | 25,733 | 202 | 93.89 | 62.78 | 3,385 |
| CIDM-BH8 | 2,939,553 | 28,704 | 193 | 93.89 | 62.78 | 3,382 |

### Sequenced *Bordetella pertussis* isolates exhibited IS*481* mediated genomic rearrangements

Three strains of *B. pertussis* were sequenced with long-read sequencing, and closure of each genome was successful. *B. pertussis* strain CIDM-BP3 and CIDM-BP5 carried intact genes of *prn* with CIDM-BP5 demonstrated to express PRN in previous work^21^. CIDM-BP2, however, contained an *IS*481 insertion at position 1598 of *prn. B. pertussis* strain CIDM-BP3 was a *ptxP1* isolate, previously described as belonging to the non-Cluster I SNP cluster with a SNP profile of 18 (SP18)^21^. Sequence alignments using the closed genome of CIDM-BP3 corroborated with the previous data^21^. Both *B. pertussis* strains CIDM-BP2 and CIDM-BP5 harboured the *ptx3* promoter and by definition^10^ were both part of SNP Cluster I. Further pairwise alignment analysis by BLASTn of CIDM-BP2 and CIDM-BP5, against key SNP positions, classified them both as part of EL1, designated SP12 and SP13, respectively, due to the SNP mutations in *gor, sphB3*, and BP3546.

The 23S ribosomal RNA was also investigated via *in silico* analysis and no mutations were discovered in any *B. pertussis* strains. This indicates that these strains are macrolide susceptible and is in line with phenotypical results for resistance against macrolides (Supplementary Material). A selection of global closed *B. pertussis* isolates (n=192) (Figure 1) and previously described^14, 15^ genomes used for structural clustering (n=472) (Figure S1) were utilised for core phylogenomic and genomic rearrangement analysis respectively. Results of both analyses revealed CIDM-BP2 aligned with more recent U.S.A and UK *ptxP3* isolates and like CIDM-BP2, the international strains contained an IS*481* inserted at position 1598 of the PRN gene. In addition, CIDM-BP2 shared the same genomic rearrangements previously observed in J247, I755, I228, I469, and I472 from Cluster-BP-12 (Figure 2).

**Figure 1:**
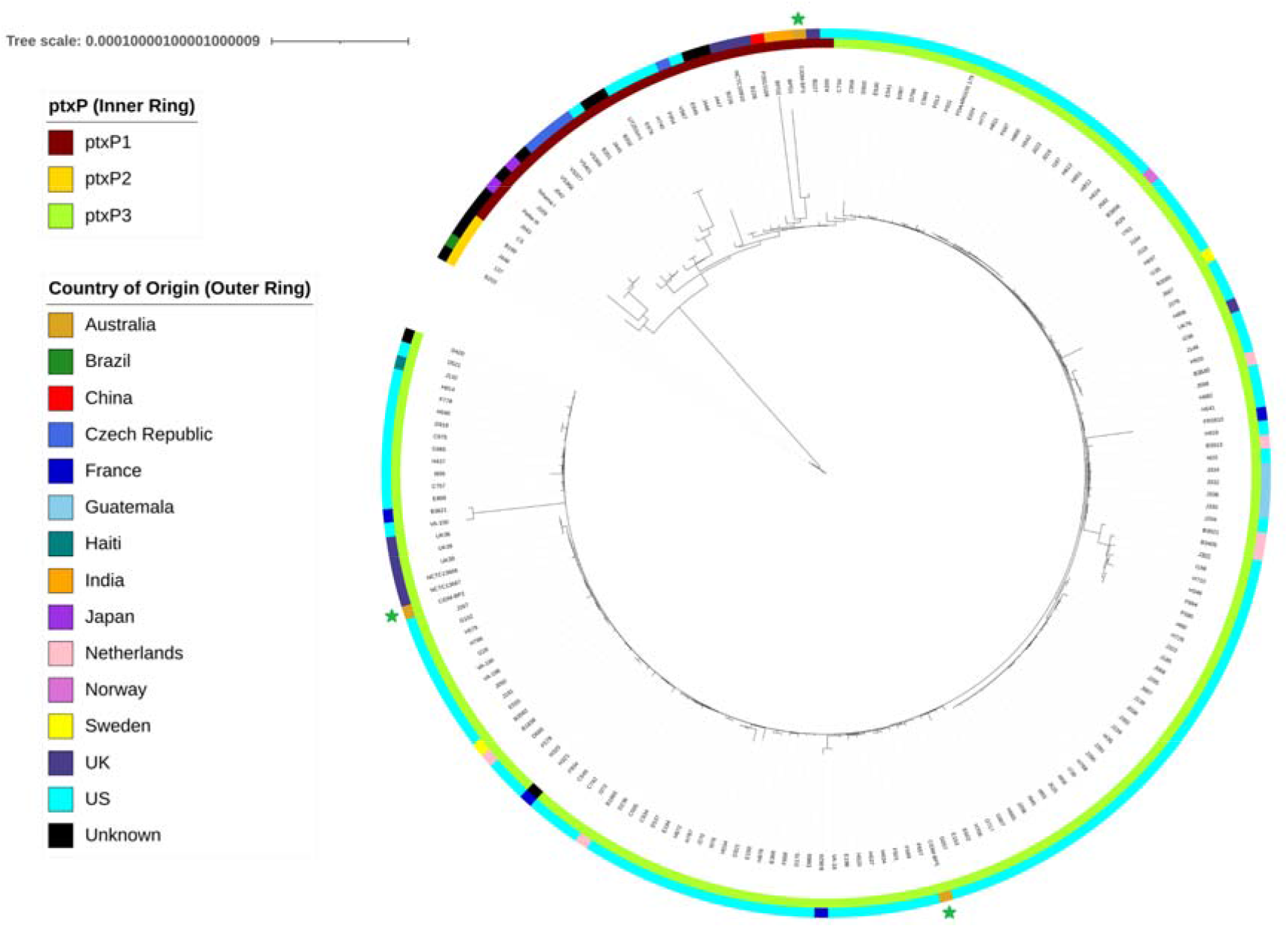
Core gene phylogeny of global closed *B. pertussis* isolates (n=192) based on 3,383 core genes using Roary. The tree shows that the *B. pertussis* isolates selected for long-read sequencing in this study (green star) represent different strains that circulate globally. The inner ring denotes *ptxP* alleles and the outer ring indicates country of origin.

**Figure 2:**
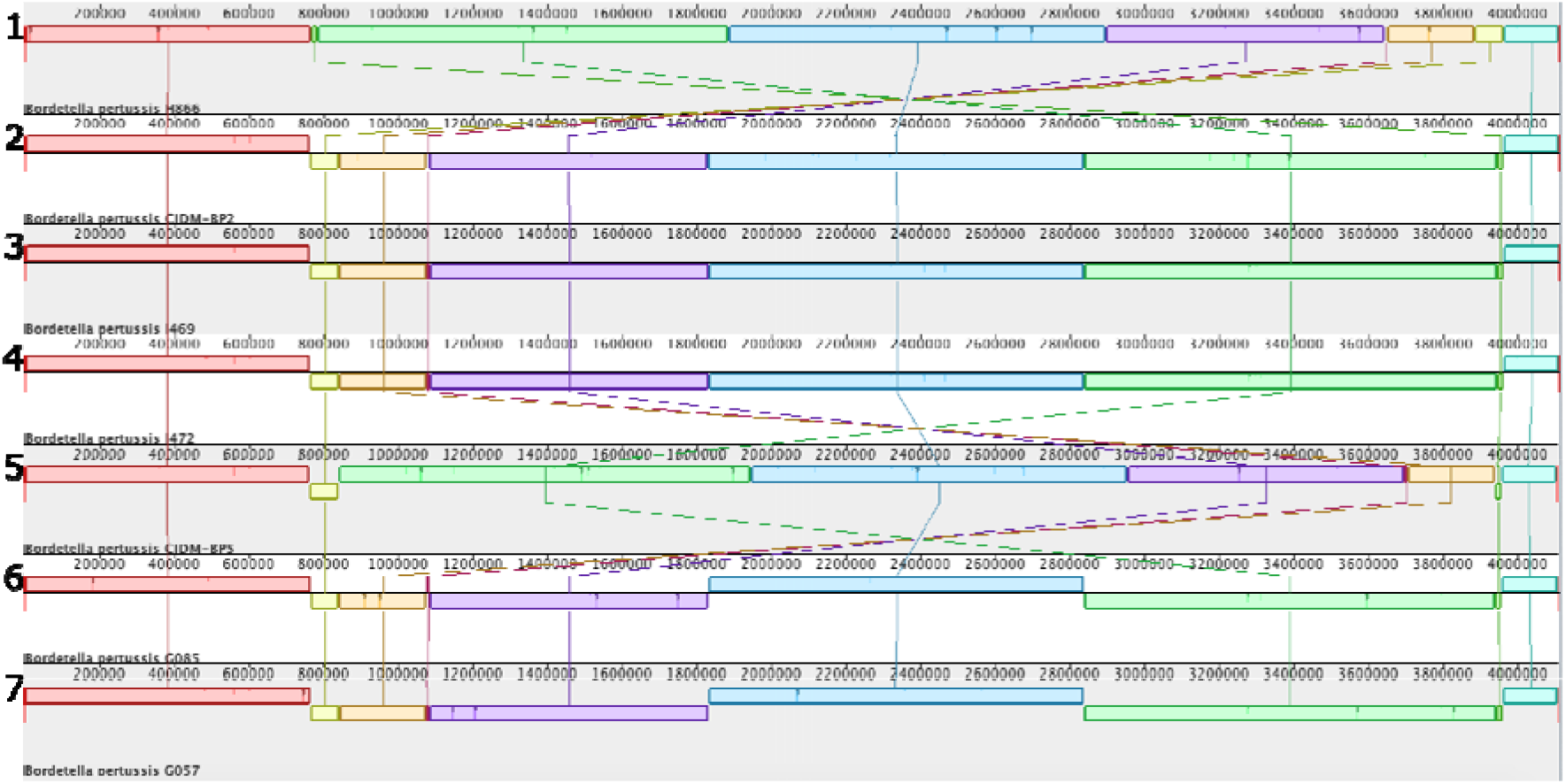
Genome structural variability comparisons of closed *B. pertussis ptxP3* genomes. Australian strains CIDM-BP2 and CIDM-BP5 are in position 2 and position 5 respectively, and are shown alongside their closest relatives by core gene phylogeny, I469 and I472 for CIDM-BP2, and G085 and G057 for CIDM-BP5 (Figure 1). H866 was used to allow a comparison with a previous study. Image generated by progressiveMauve.

Meanwhile CIDM-BP3 is closely related to *ptxP1* isolates from India (2017) and the U.K (1967) with a fully intact *prn* gene. CIDM-BP3 was observed to be more similar to strains in Cluster-BP-19 from the U.K (B226 and B228) rather than the United States (Figure S1). In terms of genomic structural organisation, CIDM-BP3 possessed the same orthologous locally collinear blocks (LCB) as H866, B226 and B228 (Figure 3) but differed from these isolates due to IS*481* mediated inversions and hence designated as a singleton. Similarly, CIDM-BP5 was also designated as a singleton due to it having a genomic organisation that was not previously documented. In addition, inversions resulted in an insertion of an *IS481* element downstream of *lysR* family transcriptional regulators in *ptxP3* isolates on two occasions, once in CIDM-BP2 and once in CIDM-BP5 (Figure 3 and Supplementary material 2).

**Figure 3:**
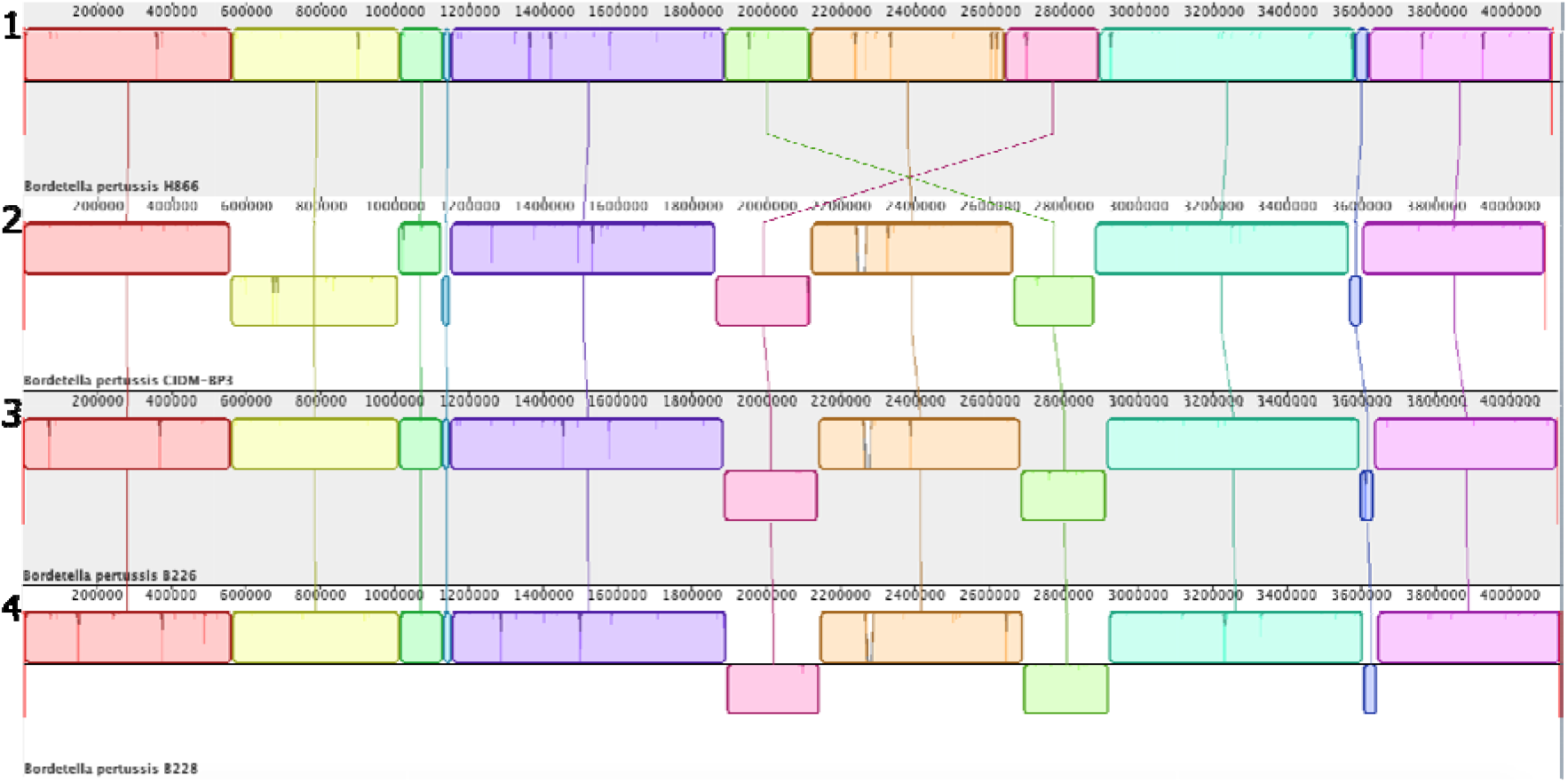
Structural rearrangement comparisons of closed *B. pertussis ptxP1* genomes, with *B. pertussis* H866 as the anchoring genome The singleton CIDM-BP3 (position 2) is featured in this figure comparing it to neighbouring isolates (B226 and B228) from the core gene phylogeny. Image generated by progressiveMauve.

**Figure 4:**
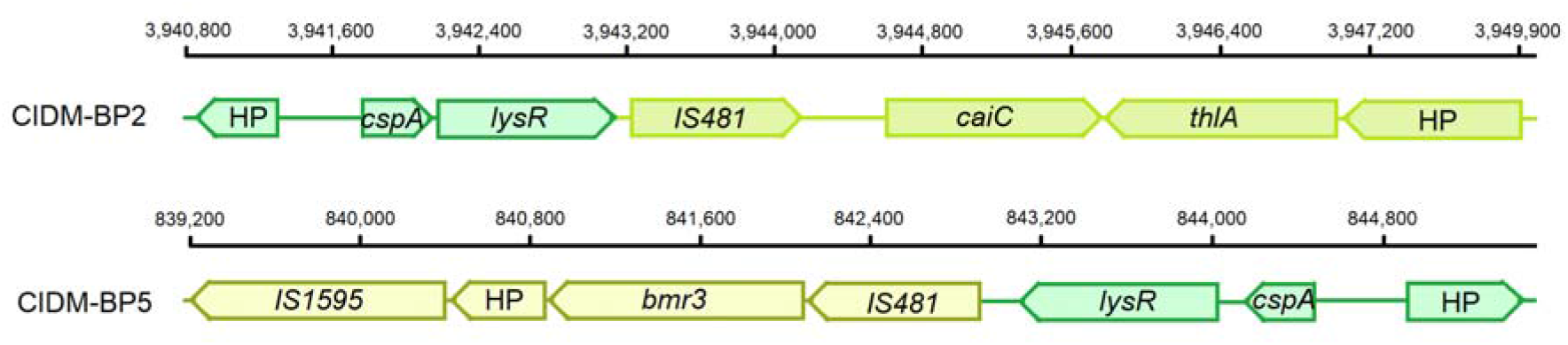
Schematic diagram of the two selected genomic rearrangement sites of CIDM-BP2 and CIDM-BP5. Represented here is the *IS481* element and three upstream and downstream flanking genes, the colours are derived from the LCB blocks in Figure 2. In both examples the *IS481* element has inserted downstream of *lysR*, as represented by the colours of the *IS481*. The flanking gene names are as follows – hypothetical protein (HP), cold shock protein (*cspA*), crotonobetaine/carnitine--CoA ligase (*caiC*), acetyl-CoA acetyltransferase (*thlA*) and Multidrug resistance protein 3 (*bmr3*).

### CIDM-BPP2 possessed a genomic structure that was previously observed

Comparison of CIDM-BPP2 to available RefSeq sequences (n=79) demonstrated that CIDM-BPP2 clustered with B160 (Figure 5). Singleton B160 (and CIDM-BP2) differs from Cluster-PP-02 by a 1,608 bp deletion in an Invasin gene (GenBank: AP019378.1). Therefore, CIDM-BPP2 forms a new cluster with B160. A loss of tandem repeats within the Invasin gene in Cluster-PP-02, resulted in the truncation of gene length, however, appears to not disrupt gene function. Examination of virulence markers of the *B. parapertussis* isolate revealed the presence of two genes that were part of a type VI secretion system (T6SS), homologous to *Pseudomonas aeruginosa* TssB1/HsiB1/VipA and TssC1/HsiC1/VipB T6SS (78% homology). Further examination showed unlike the *B. parapertussis* Bpp5 reference genome (Genbank: NC_018828.1) which contains only 7 of the 13 T6SS core components, CIDM-BPP2 and *B. parapertussis* 12822 (Genbank: NC_002928.3) contains all 13 core components, and designated T6SS Type I subtype i3 (Supplementary Table 1)

**Figure 5:**
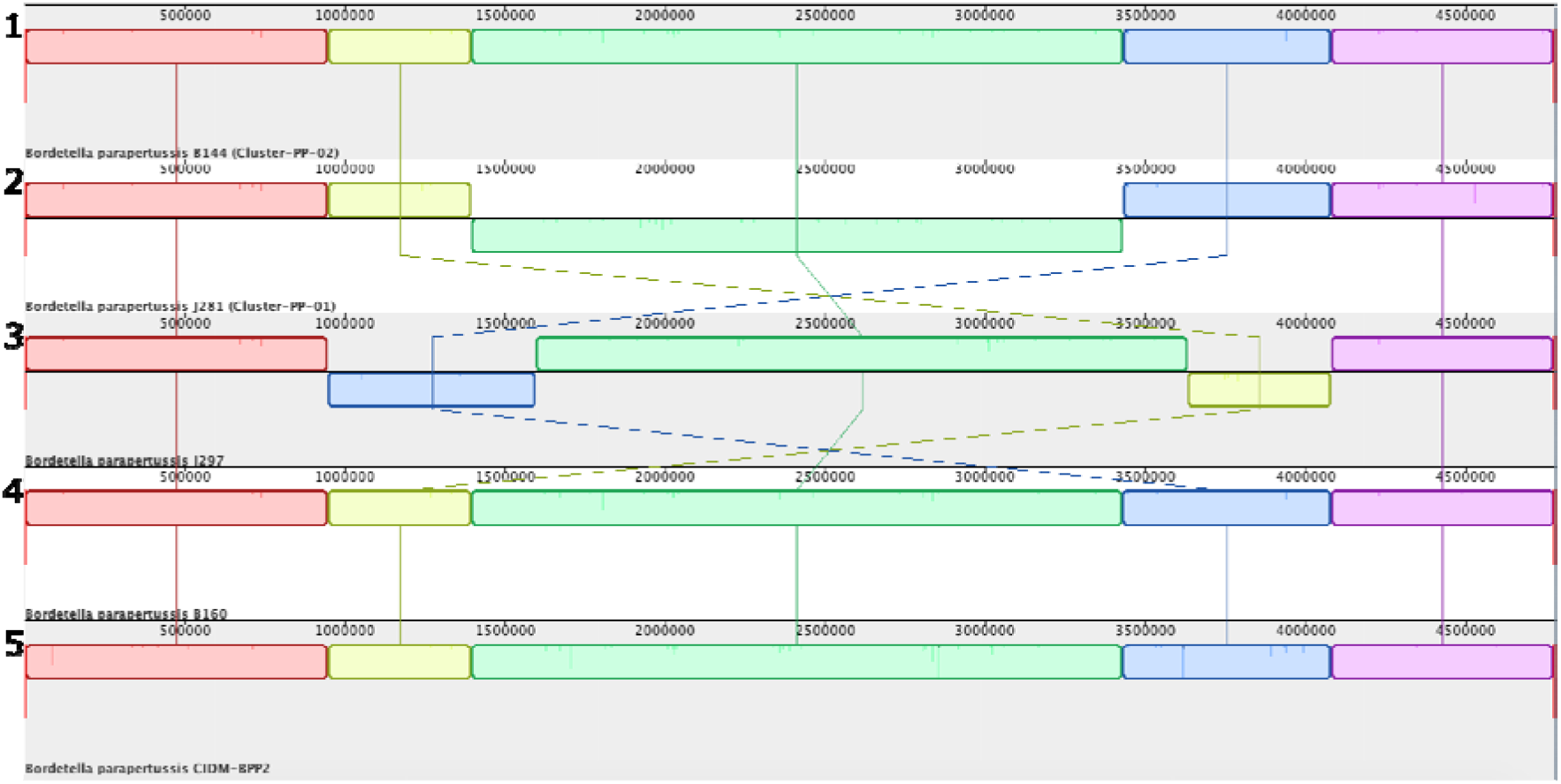
Structural rearrangement comparisons of all *B. parapertussis* structural clusters referenced by a representative isolate selected from Cluster-PP-02 (position 1). J297 (singleton) is in position 3 as featured in a previous study and B160 (singleton) is in position 4 as it was the closest strain to CIDM-BPP2 (position 5) in the core phylogenetic analysis (Figure S2). This figure shows that Cluster-PP-02 and B160 contain the same LCB orientations, however, in depth analysis reveals a 1.6kb deletion in Cluster-PP-02 isolates (genome coordinates 4,637,269 to 4,642,986). Image generated by progressiveMauve ^22^

### Global Comparison of *B. holmesii*

Phylogenetic analysis of all available *B. holmesii* isolates on RefSeq (n=84), SRA (n=10) and local (n=8), showed that CIDM-BH3 groups with Cluster-BH-01 as LCBs matched the genomic structure of the selected representative H869 (Figure 6). Like *B. pertussis* the structural rearrangements were mediated by *IS*481.

**Figure 6:**
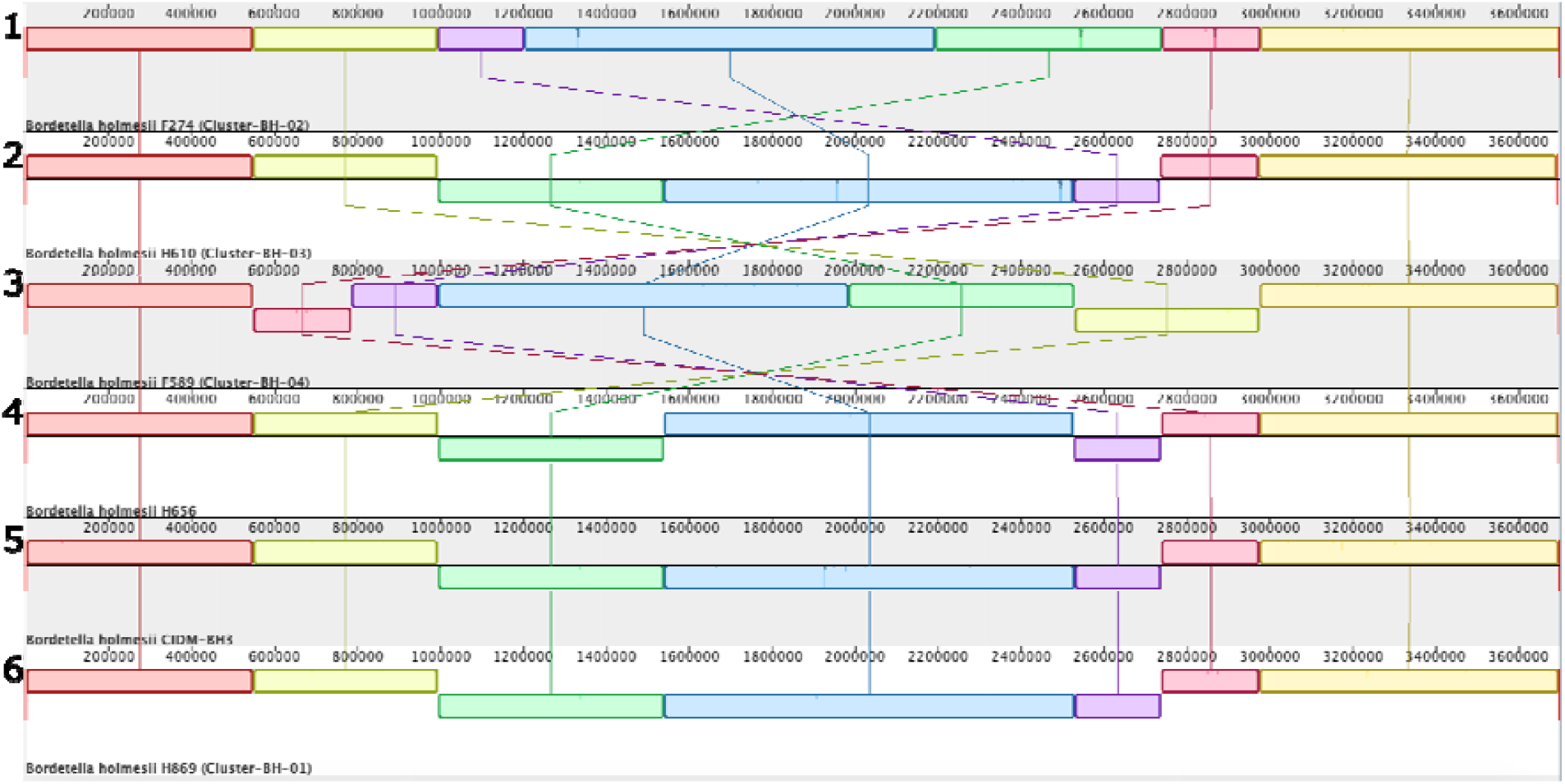
Genome structural rearrangements of *B. holmesii*. All structural clusters are featured, anchored by a representative isolate from Cluster-BH-02 (F274). CIDM-BH3 is featured in position 5, alongside its closest isolates H656 and H869 from Cluster-BH-01. Image generated by progressiveMauve^22^.

Further core genome analysis also disclosed that certain clusters display a similar absence of genes. A sub-cluster in Cluster-BH-01 is highly distinguished from other closed isolates due to a group of 53 missing genes. Half of the missing genes are IS elements, the remaining are listed in Supplementary Material 2. Whole genome screening of virulence markers using Abricate to interrogate VFDB, revealed that *B. holmesii* contains seven of the 10 homologs of the *B. pertussis bpl* locus, ranging from 76.39-83.04% identity to 91.81-98.86% coverage. *B. holmesii* lacks genes *bplG* (sugar transferase), *bplH* (glycosyl transferase) and *bplI* (lipopolysaccharide biosynthesis protein), which are genes involved in band-A lipopolysaccharide biosynthesis in *B. pertussis*^23^. Overall, *B. holmesii* lacks 87% of the virulence markers of *B. pertussis* as identified by VFDB and confirmed by BLASTn.

In total, there were 468 unique core SNP variants across Australian *B. holmesii* isolates (Supplementary Material 2). CIDM-BH1, CIDM-BH4, CIDM-BH6 and CIDM-BH7 formed a cluster (Figure 7) with 0 – 34 core SNPs between them. CIDM-BH1 and CIDM-BH4 differed from each other by 1 SNP, which is consistent as they were isolated from the same patient.

**Figure 7:**
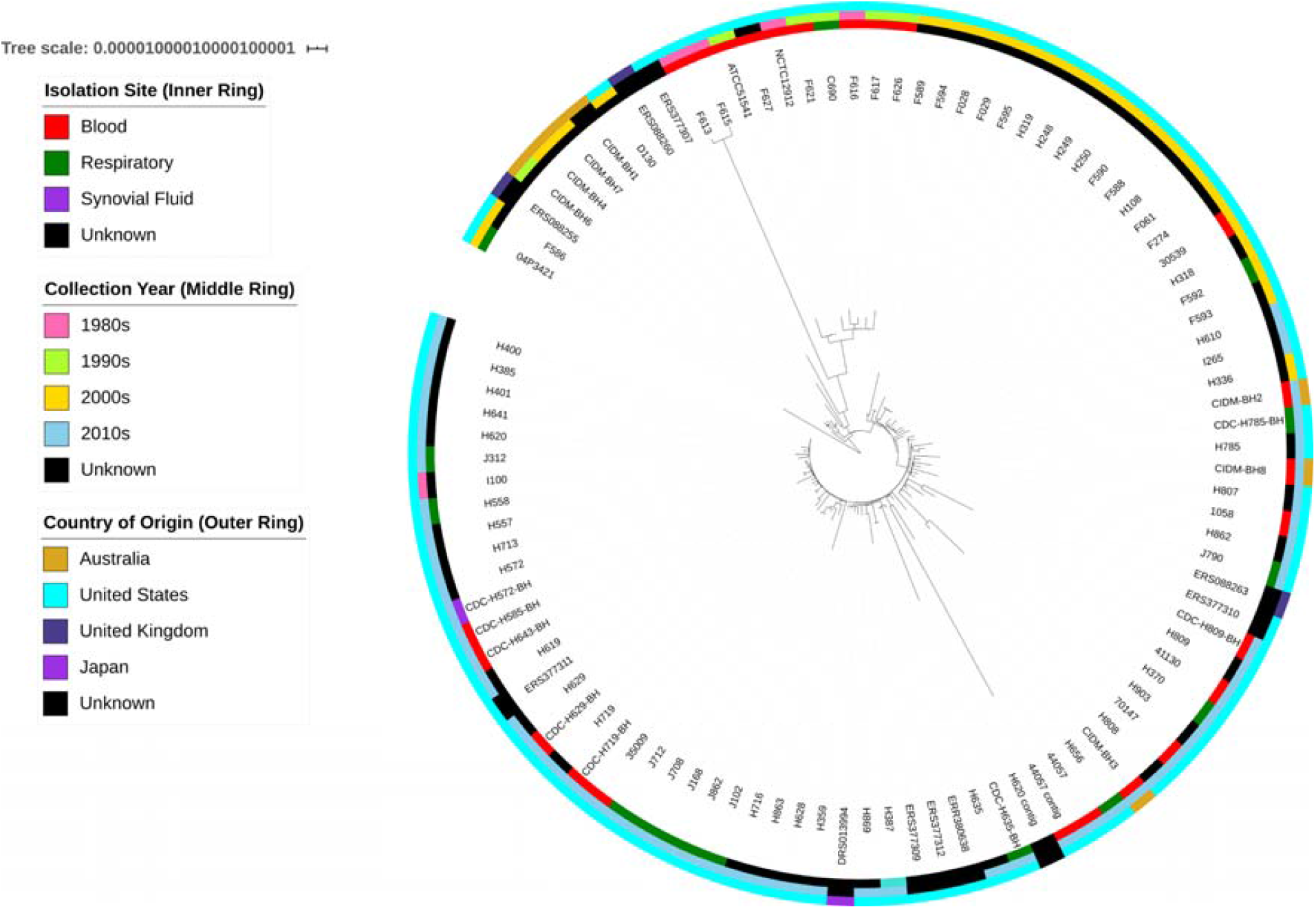
Core phylogenetic tree of all *B. holmesii* (n=101) used in the present study from NCBI SRA and RefSeq based on 2,747 core genes. Metadata is displayed in concentric circles with site of isolation (inner ring), year of collection (middle ring) and country of origin (outer ring) where available. The tree shows that Australian isolates pre-2007 are clustered together, while more recent isolates post-2012 are spread out across the globe with no specific grouping.

## Discussion

Evidence of changes in the molecular epidemiology of *B. pertussis* have been prevalent across the globe^24^. Further research into identifying forces driving this change are critical to prevent and control the re-emergence of *B. pertussis*. This study aimed to contextualise Australian *Bordetella* spp. globally. By utilising both long and short-read sequencing technologies this study generated high quality reference sequences of circulating strains of *Bordetella* spp. in Australia and provided further evidence of genomic evolution within the genus.

The *B. pertussis* genomes sequenced contained representatives of the predominating strain (*ptxP3*) in NSW with one being PRN-positive (CIDM-BP5) and one PRN-negative (CIDM-BP2), and a unique *ptxP1* PRN-positive isolate from 2015 (CIDM-BP3). *In silico* analysis confirmed that CIDM-BP3 and CIDM-BP5 each harboured an intact *prn* gene with no evidence of IS*481* insertion, supporting their phenotypic PRN production^21^. The *prn* gene in CIDM-BP2 contained an IS*481* insertion in position 1598, thus rendering the *prn* gene defective. These isolates were also classified with a previously described SNP clustering system, CIDM-BP2 and CIDM-BP5 are SNP Cluster I EL1 as SP12 and SP13 respectively, and CIDM-BP3 is a non-Cluster I SP18^21^. The isolates were also screened for known resistance mechanisms as macrolide resistant strains of *B. pertussis* have been observed in China. Macrolide resistance is the result of a 23S ribosomal RNA SNP mutation^25, 26^ and none of the study isolates exhibited genotypes consistent with resistance in this gene.

Australian *B. pertussis* isolates harbour genomic rearrangements similar to those described previously in the northern hemisphere^14, 15^. Utilising the same clustering system, CIDM-BP3 and CIDM-BP5 were found to be singletons and CIDM-BP2 was part of Cluster-BP-12. For *B. parapertussis*, CIDM-BPP2 aligned closest with Cluster-PP-02 and singleton B160. However, with a deletion of a 1.6kb region in Cluster-PP-02, it supports the addition of CIDM-BPP2 to singleton B160, and the formation of a new cluster. CIDM-BH3 was more closely related to other closed *B. holmesii* isolates from the U.S.A and clustered with Cluster-BH-01. Interestingly, core genome phylogeny can predict structural clusters, however, full confirmation of genome rearrangements is only possible with closed genome assemblies. Phylogenetic analysis on its own goes some way to predict the structural cluster of draft genomes, however full confirmation requires long-read sequencing at a cost that is significantly higher than short-read high throughput sequencing.

The genomic inversions reported previously^14, 15^ here, can be mediated by insertion sequences, primarily *IS481* in *B. pertussis* and *B. holmesii*, and *IS1001* in *B. parapertussis*. As in previous work, the study found that *IS481* and *IS1001* flanked all ends of all identified rearrangements. These IS are present in multiple copies in all *Bordetella* genomes and can number at over 200 copies in *B. pertussis*. The impact of these IS elements requires further investigation as previous studies have indicated that IS elements can upregulate upstream and downstream flanking gene expression in *B. pertussis*^27^. This lends to the hypothesis that specific structural clusters could be more virulent than others^27^, as in some cases, *IS481* was introduced in front of transcriptional regulators post-inversion. The hypothesis can be addressed using RNA-Seq to resolve the transcriptome of strains from different structural clusters and comparing the impact of gene expression due to *IS481*.

A deeper core genome analysis into *B. holmesii* isolates in NSW showed that between 1999 and 2012, isolates were part of a specific cluster, differing by up to 32 SNPs only. More recent isolates (post-2012), however, differ from pre-2012 isolates by more than 72 SNPs indicating a potential change in circulating strains. Although, the small number of *B. holmesii* isolates in NSW makes a solid conclusion hard to draw. Given the more recent discovery of *B. holmesii* causing pertussis-like symptoms, a culture-independent approach to sequencing can be applied to *B. holmesii* positive respiratory samples to potentially close this gap in knowledge of the molecular epidemiology of *B. holmesii* in Australia.

Virulence marker investigations across the *Bordetella* spp. revealed that various virulence-associated genes of *B. pertusiss*, were not present on the genome of *B. parapertussis* or *B. holmesii*. In *B. parapertussis* CIDM-BPP2, all components of a T6SS were discovered, homologous to the T6SS found in *P. aeruginosa* and *V. cholerae*. This T6SS has also been discovered in *B. bronchiseptica* RB50, and whether the T6SS in *B. parapertussis* remains functional will have to be further investigated. *B. parapertussis* carries all the same virulence genes as *B. pertussis* with varying mutations including the *ptx-ptl* operon, however, as previously demonstrated, this operon in *B. parapertussis* is dysfunctional^28^. In comparison, *B. holmesii* lacks a large majority (87%) of the virulence markers found in *B. pertussis*, these include the vaccine antigens of *B. pertussis*.

In conclusion, the study applied structural clustering on closed genomes of the *Bordetella* spp. to determine where NSW isolates sit on a global scale. This study also presents the first five high-quality Southern hemisphere closed genomes of *Bordetella* spp. These reference genomes are crucial for future public health investigations of this significant group of respiratory pathogens given the level of genomic rearrangement displayed by all species investigated in this study. While this study demonstrates further evidence of genomic evolution of the *Bordetella* species, dwindling isolate numbers due to diagnostic testing changes hamper the capability to continue monitoring *Bordetella* spp.

## Methods

### Strain selection and culture conditions

Chosen strains (Table 3) were stored at the Centre of Infectious Diseases and Microbiology Laboratory Services (CIDMLS) Identification Laboratory were cultured on Charcoal Blood Agar (*B. pertussis* and *B. parapertussis*) without Cephalexin (CBA) or Horse Blood Agar (HBA) (*B. holmesii*) for 7 days at 37°C.

**Table 3:** *Bordetella* spp. strains selected for both Nanopore and Illumina sequencing

| Strains | Isolation Year | Isolation Site | Sequencing |
| --- | --- | --- | --- |
| *Bordetella pertussis* |  |  |  |
| CIDM-BP2 | 2011 | Respiratory | Nanopore |
| CIDM-BP3† | 2015 | Respiratory | Nanopore |
| CIDM-BP5 | 2008 | Respiratory | Nanopore |
| *Bordetella parapertussis* |  |  |  |
| CIDM-BPP2 | 1993 | Respiratory | Nanopore |
| *Bordetella holmesii* |  |  |  |
| CIDM-BH1 | 2000 | Blood | Illumina |
| CIDM-BH2 | 2014 | Blood | Illumina |
| CIDM-BH3 | 2016 | Blood | Nanopore |
| CIDM-BH4 | 2000 | Blood | Illumina |
| CIDM-BH6 | 1999 | Blood | Illumina |
| CIDM-BH7 | 2007 | Unknown | Illumina |
| CIDM-BH8 | 2012 | Blood | Illumina |

†This isolate has been published previously as L2263/BP7<sup>21</sup>

### DNA Extraction and Quality Control Metrics

Two forms of DNA extraction were employed in this study, as not all strains were sequenced by Nanopore sequencing on the MinION platform (Table 3). DNA was extracted from isolates for long read sequencing (Nanopore) using the DNeasy^®^ UltraClean^®^ Microbial Kit (Qiagen, Germany) with modifications. Briefly, two loopfuls of culture were picked and inoculated into the Powerbead tube provided containing both Powerbead solution and solution SL. Cells were lysed mechanically for 2 minutes and subsequent steps were performed according to Manufacturer’s instructions. DNA integrity was assessed by running the extracted DNA on a 0.6% (w/v) agarose gel for 3 hours at 75V in 1X TBE sized with a Quick-Load^®^ 1 kb Extend DNA ladder (New England Biosciences, USA) to ensure that majority of the fragment sizes were > 30 kb.

Isolates for short read sequencing (Illumina) were extracted with the DNeasy Blood and Tissue kit (QIAGEN, Germany) with modifications. Two colonies were picked from the culture plates and inoculated directly into the buffer supplied in the kit and Proteinase K. The cells were lysed at 56°C on a heater-shaker for 3 hours, and subsequently extracted following the manufacturer’s instructions.

All extracted DNA were quality assessed spectrophotometrically (AllSheng, China) and DNA quantity for long read sequencing was measured on a Qubit^™^ 2.0 Fluorimeter using the dsDNA BR Assay Kit (ThermoFisher Scientific, USA). DNA quantity for short read sequencing was measured using Quant-iT^™^ PicoGreen^™^ dsDNA Assay Kit (ThermoFisher Scientific, USA). DNA was stored in −20°C until library preparation.

### Library Preparation and Sequencing

All extracted DNA was prepared with the Illumina NexteraXT DNA Library Preparation Kit v2.5 (Illumina, USA) and sequenced on the Illumina NextSeq 500.

Libraries for isolates that were sequenced on the Nanopore was prepared using the Rapid barcoding Kit SQK-RBK 004 (Oxford Nanopore, U.K) using > 1,000 ng of input DNA. Apart from the change in input amount, all subsequent manufacturer’s instructions were followed. DNA was sequenced on a FLO-MIN106 flow cell for 24-48 hours without basecalling.

### Bioinformatics

#### Base calling, assembly and annotation

Long read base calling in high accuracy mode and demultiplexing was performed on Guppy (v 2.4.5) on a GPU Amazon Web Service instance (c5.Metal). Demultiplexed reads were then *de novo* assembled with Flye (v 2.7b)^29^ with the “--asm-coverage” parameter set to 30 and an expected genome size of 4.0 Mb. Following long-read assembly, the sequence was corrected with Racon (v 1.3.1)^30^ four times, and Medaka (v 0.11.5)^30^ twice. The assembly was then polished with corresponding Illumina reads using Pilon (v 1.23)^31^ and repeated until there were no more changes. The assembly was subsequently reordered so that the open reading frame of *dnaA* was in the positive strand.

All short read sequences were assembled with SPAdes (v 3.12.0)^32^, then annotated with Prokka (v 1.12)^33^ and Barnapp (v 0.6) (https://github.com/tseemann/barrnap), then scanned for virulence factors (VFDB)^34^ and resistance markers (CARD)^35^ with ABRicate (v 0.9.8) (https://github.com/tseemann/abricate). The type VI secretions system analysis was performed by SecReT6^36^.

### Phylogenetic analysis

To determine where the Australian *Bordetella* spp. isolates sat globally, all closed RefSeq genomes were downloaded. Raw reads from SRA were also utilised and assembled with SPAdes (v 3.12.0)^32^. All assemblies and closed sequences were annotated with Prokka (v 1.12)^33^ and Barnapp (v 0.6) (https://github.com/tseemann/barrnap).

For *B. pertussis*, a random selection of RefSeq sequences were extracted based on country and year of isolation (n=189) In addition, previously described closed genomes^14^ were downloaded from RefSeq (n=472) and reanalysed with Australian *B. pertussis* isolates. SNP clusters were determined by BLASTn^37^ with a custom database of SNPs positions from a previously published study^10, 11^. *B. holmesii* analysis was performed with all available RefSeq (n=84) and SRA sequences (n=10). SNP analysis of *B. holmesii* was performed by Snippy (v 4.1.0) and SNP-dist (v 0.6). For *B. parapertussis*, only closed sequences featured in Weigand et al were used for analysis. A list of all strains used in the study are in Supplementary Material 1.

For each *Bordetella* spp. generated GFF files from Prokka were used for phylogenetic analysis with Roary (v 3.1.2)^38^, and the tree was drawn with IQ-TREE (v 1.6.7)^39^ and ModelFinder^40^, with the best model being TVM+F+I. The core phylogenetic tree was visualised with iTOL (v 5.7)^41^. Utilising the phylogenetic results, strains closest to the Australian isolates were extracted for structural rearrangement analysis with progressiveMauve (v 2.4.0)^22^, the same cut-offs applied by Weigand et al^14^ were applied in the study.

Genome sequencing data has been published on NCBI under Bioproject: PRJNA695314

